# Structural genomic variation and migratory behavior in wild songbirds

**DOI:** 10.1101/2023.04.24.538030

**Authors:** Kira E. Delmore, Benjamin M. Van Doren, Kristian Ullrich, Teja Curk, Henk P. van der Jeugd, Miriam Liedvogel

## Abstract

Structural variants (SVs) are a major source of genetic variation, but accurate descriptions in natural populations and connections with phenotypic traits remain scarce. We integrated advances in genomic sequencing and animal tracking to begin filling this knowledge gap in the Eurasian blackcap. Specifically, we (i) characterized the genome-wide distribution, frequency and overall fitness effects of SVs using haplotype-resolved assemblies for 79 birds, and (ii) used these SVs to study the genetics of seasonal migration. We detected >15K SVs. Many SVs overlapped repetitive regions and exhibited evidence of purifying selection suggesting they have overall deleterious effects on fitness. We used estimates of genomic differentiation to identify SVs exhibiting evidence of selection in blackcaps with different migratory strategies. Insertions and deletions dominated these SVs and were associated with genes that are either directly (e.g., regulatory motifs that maintain circadian rhythms) or indirectly (e.g., through immune response) related to migration. We also broke migration down into individual traits (direction, distance and timing) using existing tracking data and tested if genetic variation at the SVs we identified could account for phenotypic variation at these traits. This was only the case for one trait – direction – and one specific SV (a deletion on chromosome 27) accounted for much of this variation. Our results highlight the evolutionary importance of SVs in natural populations and provide insight into the genetic basis of seasonal migration.

## Introduction

Structural variants (SVs) include duplications, deletions, transpositions and inversions. Existing research suggests that these variants represent a major source of genetic variation and could have important fitness consequences (1–3). For example, SVs can have deleterious effects on fitness, disrupting functional features of the genome (e.g., exons) and/or suppressing recombination (4–6). Suppressed recombination can lower effective population size at the local genomic level, reducing the efficiency of purifying selection. There is growing evidence SVs could also have the opposite effect, facilitating adaptation and speciation (7–9). For example, reductions in recombination can also allow co-adapted alleles at separate loci to segregate together. This co-segregation can shelter co-adapted alleles that underlie adaptive phenotypic traits. If these phenotypic traits are important for maintaining reproductive isolation and gene flow with other populations is occurring, co-segregation could also facilitate speciation (10–12).

Despite their potential fitness effects, data on the genome-wide distribution of SVs and their frequency within populations is lacking. This dearth of knowledge is especially true in natural populations of non-model organisms and is related in large part to technological limitations. Specifically, advances in sequencing technology have made it possible to obtain genome-wide data from non-model organisms, but existing work is often limited to short (∼150 bp) sequencing reads. SVs are often larger than these reads and can be highly repetitive, making them difficult to assemble (13–15). A complete understanding of SVs will require contiguous genomes from multiple individuals where repetitive regions of the genome have been assembled accurately (16,17). Linked reads are one technology that can help meet this need. They use molecular barcoding to preserve long range sequencing information. Here we used this technology to identify SVs in a natural population of European blackcap. We have matching data on the migratory behavior of each bird, allowing us to gain inference into the genetics of seasonal migration as well.

Seasonal migration is the yearly long-distance movement of individuals between their breeding and wintering grounds. Successful migration requires the integration of several behavioral, physiological and morphological traits (18,19). Decades of research has shown that there is a genetic basis to many of these traits, but the actual identity of genes underlying them remains largely unknown. Genes controlling the circadian clock have been linked to some traits, but unbiased, genome-wide studies have only recently been applied to this question (20–23). Most genome-wide studies are limited to population-level comparisons, estimating genomic differentiation between populations that differ in one or more migratory traits (24–28). These comparisons are valuable for identifying genomic regions under positive selection (i.e., areas of elevated differentiation), but caution is needed when interpreting their results as other processes can also elevate differentiation, including background selection and selection unrelated to the trait of interest (29–31). A complete understanding of migration genetics will require complementary work at the individual-level, including genome-wide association studies (GWAS) connecting specific genomic regions to individual migratory traits. GWAS will not only tell us about the genetics of individual migratory traits, but will also help us understand how these traits are integrated at the molecular level. Given their potential to shelter co-adapted alleles, SVs may be important for this integration. Indeed, there is already evidence that SVs underlie migratory traits in two avian systems; separate inversions on chromosome 1 underlie migratory orientation in willow warblers and wing shape in common quails (27,32–34).

The Eurasian blackcap is found throughout much of Europe, northern Africa and central Asia (Fig 1). This species exhibits considerable variation in migratory behavior – resident and migratory populations exist and among migrants, three main orientations have been described (northwest [NW], southwest [SW] and southeast [SE] on fall migration)(35,36)(Fig 1). SE and SW migrants form a migratory divide in central Europe. Birds at the center of this divide orient in intermediate, southern (S) directions (36). Additional differences in the distance, timing, speed and duration of both fall and spring migration have also been documented in migrants (36)(Fig 1). Researchers have capitalized on variation in the migratory behavior of blackcaps to study the genetics of migration for decades, including experimental and quantitative genetics approaches showing there is a strong genetic basis to migratory traits (37–39). Genetic surveys indicate that this variation arose recently and has not resulted in substantial, genome-wide differentiation (25,40,41). These genetic surveys include a recent study that used whole genome resequencing data to identify eight small genomic regions under positive selection in migrants that orient in different directions (four, three and one in the NW, SE and SW groups, respectively). The former study was limited to single nucleotide polymorphisms (SNPs) and population-level comparisons using individually resequenced birds distant from the contact zone with population averaged phenotypes (25).

**Figure 1.**
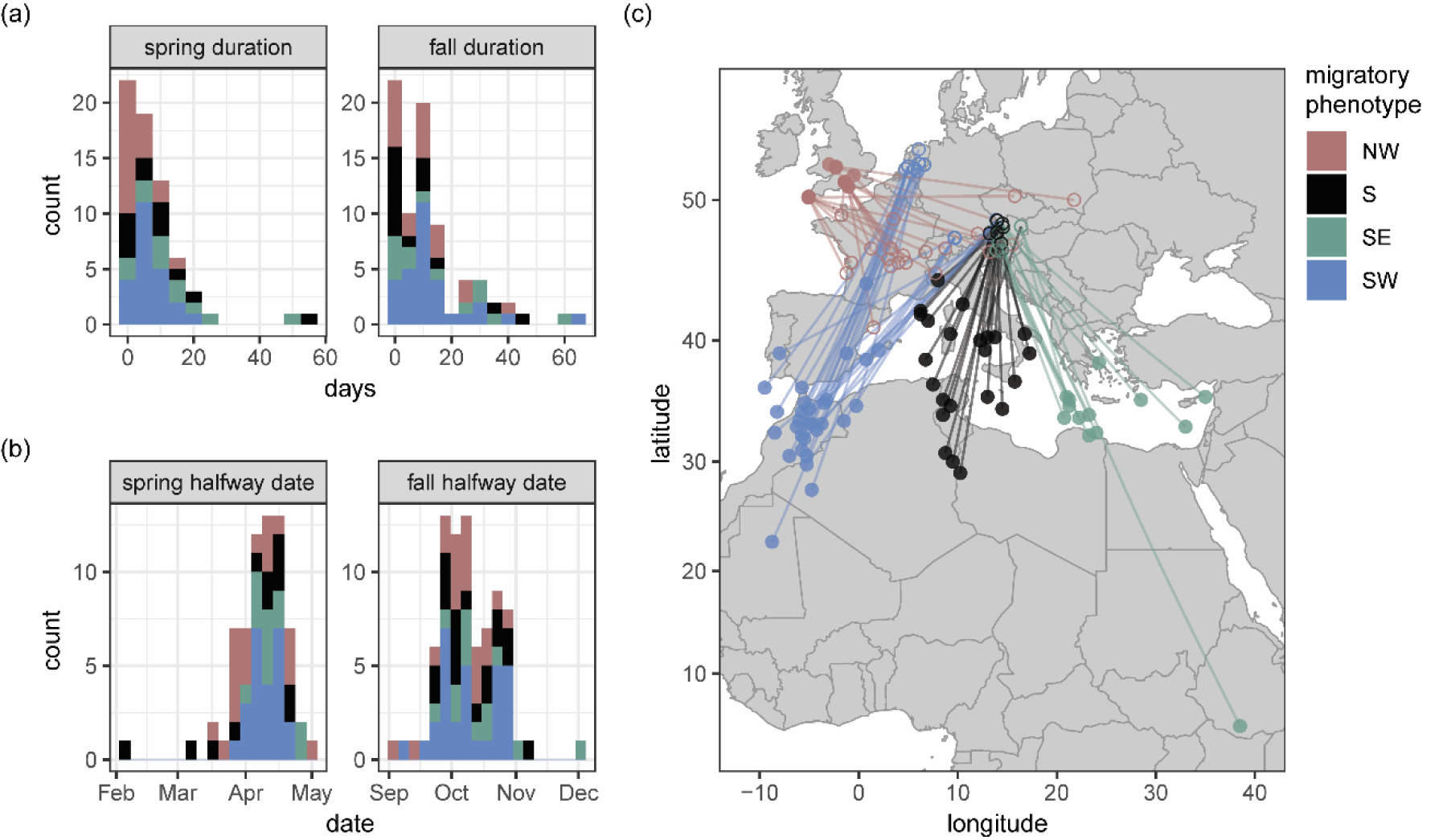
Migratory behavior of Eurasian blackcaps. Timing of (a) spring and (b) fall migration and (c) map showing wintering and breeding locations.

Here, we used linked reads to generate haplotype-resolved *de novo* assemblies for 79 individual blackcaps with individual based phenotype characterization, including NW, SW and SE migrants and individuals from the migratory divide in central Europe (Fig 1). We called SVs using these *de novo* assemblies and had three main objectives: (1) characterize SVs in natural populations, including their genome-wide distribution and overall fitness effects, (2) test for greater genome-wide population differentiation than former genetic surveys, and (3) use complementary population- and individual-level analyses to study the genetics of seasonal migration, including (a) local estimates of genomic differentiation to identify regions under selection and (b) GWAS to test if these regions are linked to specific migratory traits. All of the individuals used in the present study were tracked with light-level geolocators (36). Accordingly, we have individual-level phenotype data to run GWAS on multiple migratory traits, including direction, distance, the location of wintering grounds and both the duration and timing of fall and spring migration.

## Results and Discussion

We constructed *de novo* genome assemblies for 79 blackcaps using 10X Genomics linked-read technology (Table S1). Final assemblies averaged 999.6 Mb in size and included an average of 1,710 scaffolds. Average scaffold and contig N50 sizes were 11 Mb and 110 kb, respectively, and >91% of the universally conserved single-copy benchmark (BUSCO) genes were present in these assemblies. Given considerable uncertainty associated with calling SVs (42), we used these assemblies and three separate pipelines to genotype birds, limiting our analysis to SVs called in two or more pipelines. Following these criteria, we identified between 9,246 and 12,585 SVs per individual. After merging data from all individuals and filtering out variants with minor allele frequencies <0.05, our final dataset comprised 15,764 SVs.

### Characterizing SVs and examining their overall fitness effects

We started our analysis by examining the distribution of SVs across the genome, counting the number of SVs in non-overlapping windows of 200 kb. We found an average of three variants per window, with some windows harboring much larger numbers of SVs, especially on microchromosomes, where densities exceeded 15 SVs/window (Fig 2a).

**Figure 2.**
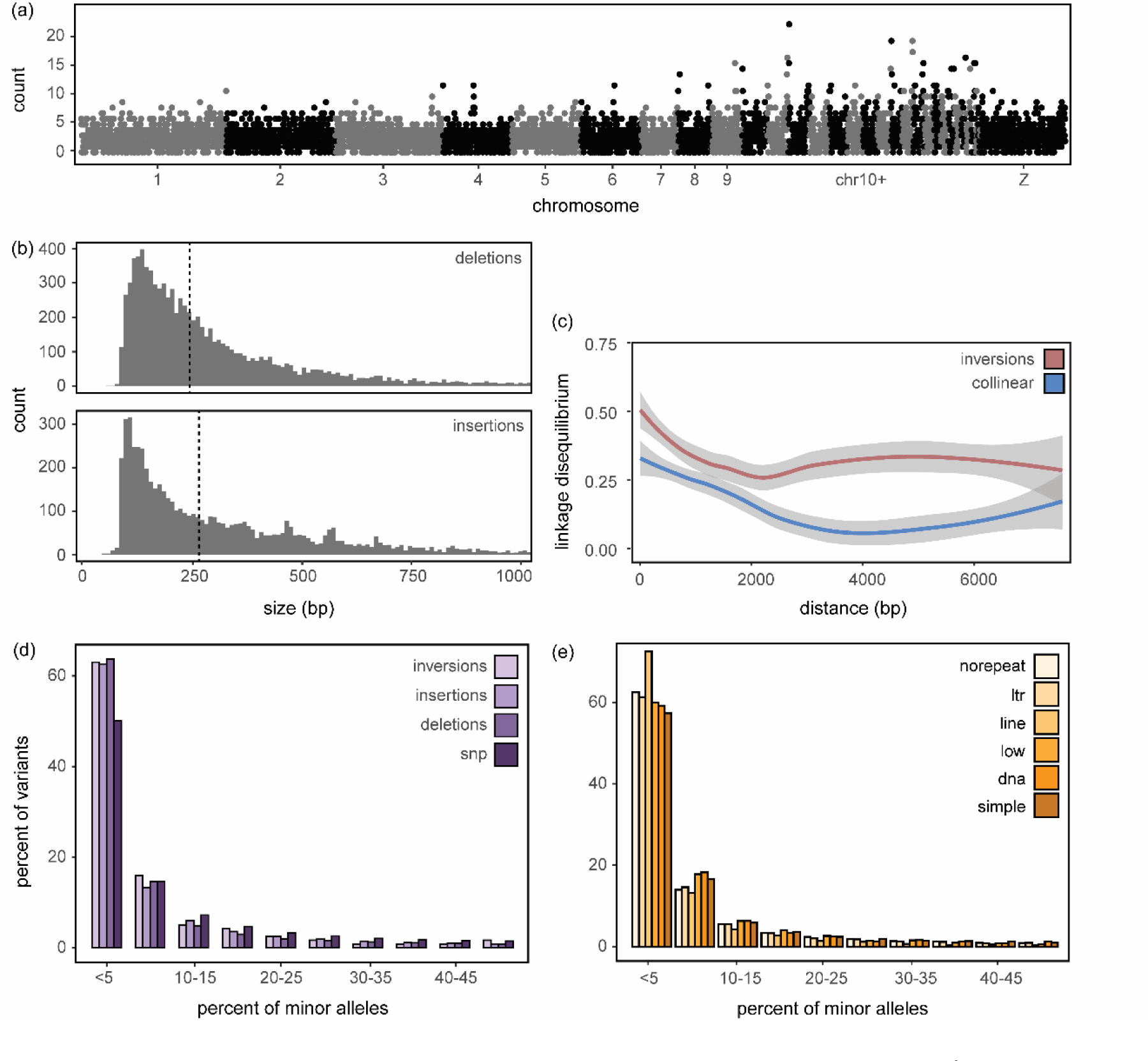
Structural variants and their fitness effects. (a) Density (number/200 kb window) along the genome and (b) size distributions for deletions and insertions including dotted line for median sizes. (c) Average decay of linkage disequilibrium as a function of physical distance in major and minor inversion homozygotes and collinear regions. Loess curves and their standard errors are shown. Allele frequency spectrums (AFS) for (d) each type of structural variant and (e) those overlapping different repeat elements in the genome.

Deletions were the most numerous type of SV (n=9341), followed by insertions (n=6393), inversions (n=24), tandem duplications (n=4) and translocations (n=1). There was a bias towards smaller SVs among deletions and insertions, with median sizes of 245 and 265 bp in each class, respectively (size range of 58 – 65285 bp for deletions and 52 – 15209 bp for insertions; Fig 2b). The tandem duplications were a similar size (median 272 bp; range 144 – 357 bp), but the median size of inversions was approximately 10 times larger (2485 bp; range 365 – 7774 bp). The size of the single translocation was 10,456 bp.

We used minor allele frequency spectra (AFS) to examine the overall fitness effects of SVs using all individuals, starting with a comparison of variant types and including an AFS for SNPs called using the same sequencing data for comparison. AFS of all SV types were skewed towards rare alleles when compared to the AFS for SNPs (Fig 2d), indicating that alleles at SVs segregate at lower frequencies than alleles at SNPs and are under stronger purifying selection (i.e., are more deleterious than alleles at SNPs). This finding is in line with theoretical predictions that SVs are often deleterious as well as results from other species using a similar approach to quantify overall fitness effects associated with SVs (e.g., *Drosophila*, (43); cacao trees, (6); European crows, (13)). Population structure can also skew AFS, but previous genomic work on the blackcap and results below reveal limited structure in this species; the most distinct variation across its distribution range coming from resident versus migratory populations but variation between populations with distinct migratory strategies is very low (25,40,41).

SVs frequently occurred in repetitive regions of the genome. We annotated SVs using a repeat library manually curated for the Eurasian blackcap (44,45). Thirty-seven percent of the variants overlapped one or more repetitive elements in this library. Close to half of the repeats that overlapped these variants were simple repeats (47.8 %); one quarter overlapped LTR retrotransposons (25.5 %). The rest were LINE/CR1 retrotransposons (17.5 %), low complexity repeats (8.2 %) and to a much smaller degree DNA transposons (0.79 %) and SINE retrotransposons (0.24 %; Fig 2e). The AFS for LINE/CR1 retrotransposons was skewed towards rare alleles suggesting they have the strongest deleterious effects. Comparable proportions of repeat elements were reported in other songbird genomes, suggesting that transposable elements (especially LTR and LINE/CR1) are highly active in this group of organisms (*Ficedula* flycatchers, (44); European crows, (13)).

Only a small number of inversions were identified in our dataset, but they exhibited molecular signatures expected of this variant type. For example, recombination between major and minor arrangements was not observed at these inversions. We estimated linkage disequilibrium (LD) in inversions and a control set of colinear regions (same sizes and number). As expected, LD dropped off rapidly in colinear regions (at ∼2000 bp). However, this was not the case in inversions, with SVs continuing to exhibit elevated levels of LD as distances between variants increased beyond 2000 bp (4000 bp and beyond) (Fig 2c).

### Genome-wide levels of population differentiation at SVs

Consistent with previous genetic surveys using molecular tools to characterize population structure of European blackcaps, we found little evidence for population structure using SVs. Previous studies using marker based approaches, such as mitochondrial haplotypes, microsatellites, but also genome wide SNP based approaches clearly show that population structure among medium-distance migrants with distinct migratory orientation is very low (25,40,41). We assigned birds from the present study to NW, SW, SE or intermediate (S) groups using their vector between breeding and wintering locations to characterize autumn migratory direction (Table S1) and tested if we could recover population structure using SVs. Specifically, we summarized genetic variation at SVs using a principal component analysis (PCA). We limited this analysis to autosomal chromosomes because our dataset comprises both males and females and the sex chromosomes accounted for a large amount of documented variation (Fig S1a). Once the sex chromosomes were excluded, only the first PC was significant (p=0.0033, eigenvalue=1.17). Birds did not cluster based on breeding or wintering location (Fig S1b). This lack of structure was true even when contrasting birds with breeding locations furthest away from the contact zone (e.g., the Netherlands vs. Austria) (Fig S1c). Combined with previous genetic surveys, these results indicate that differences in migration do not generate strong genome-wide differentiation at SVs or any other genetic marker examined in the Eurasian blackcap thus far.

### Local genomic patterns of differentiation

Low levels of genetic differentiation between populations that exhibit distinct differences in phenotypic traits are ideal for work on the genetic basis of phenotypic traits, as genomic regions that underlie these traits should standout against the backdrop of limited differentiation. Accordingly, we used a series of analyses aimed at identifying genomic regions that underlie migratory traits in blackcaps. We started with Population Branch Statistics (PBS) (47), an *F_ST_* based statistic that can be applied to comparisons with more than two groups and identifies allele frequency differences specific to each group. We limited our analysis to NW, SW and SE migrants, excluding birds exhibiting intermediate (S) orientations to contrast the most extreme phenotypes. Among SW and SE migrant, we limited our analysis to birds that bred in a geographically confined area across the migratory divide in Austria (i.e., excluded birds breeding in the Netherlands) to minimize any potential confounding effects of even small amounts of population structure.

PBS was lowest for SW birds, and very few variants stood out against baseline levels in this group (average PBS in SW = 0.009; 0.013 in both NW and SE) (Fig 3). Both NW and SE migrating birds had several SVs that stood out against baseline levels of PBS (Fig 3). Variants exhibiting extreme values of PBS may be under positive selection and important for encoding variation in migratory behavior of these birds. Accordingly, we extracted genes that overlapped SVs in the top 5% of the PBS distribution for each group (PBS > 0.06 [SW], 0.10 [NW], and 0.10 [SE]; 172 [SW], 155 [NW], and 162 [SE] genes, respectively) and ran a functional enrichment analysis, looking for gene ontology (GO) terms, biological pathways and regulatory motifs that were overrepresented in these genes. Regulatory motifs are sequences of DNA that are bound by transcription factors, one of which (Ebox) was enriched for in the NW migrants. The Ebox motif is often bound by basic helix-loop-helix (bHLH) transcription factors and is of significance for migration as several genes that regulate the circadian clock are bHLH transcription factors that bind Ebox motifs (48). In a response to changes in photoperiod, the circadian clock entrains circadian (possibly also circannual) rhythms as well as initiates migratory behavior (49,50). Interestingly, we documented similar enrichment in our previous genomic survey of blackcaps, using SNPs and PBS to identify genomic regions under selection in the same migratory phenotypes (25). No functional enrichment was found in the SW migrants.

**Figure 3.**
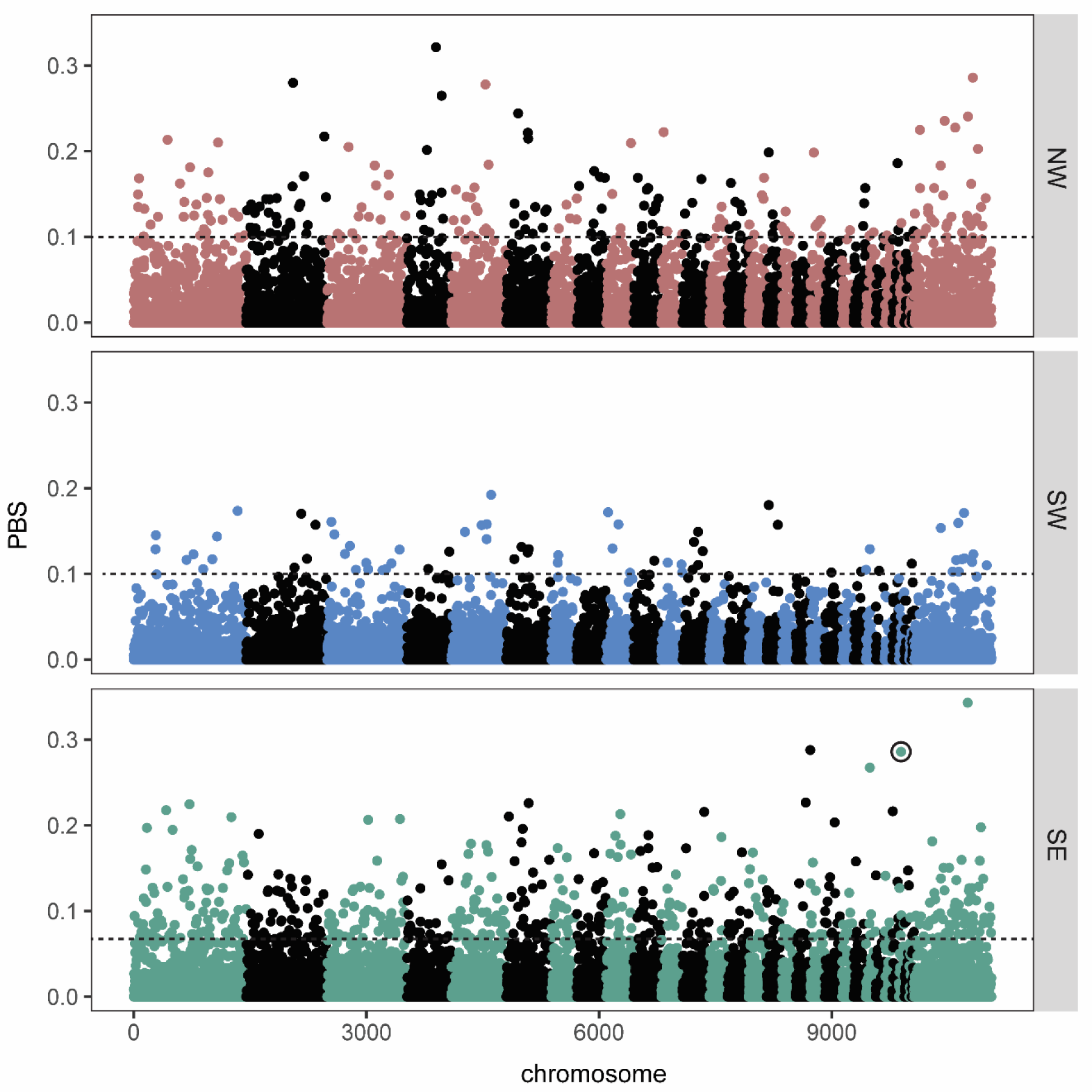
Signatures of selection. Population branch statistics (PBS) estimated for birds that migrated along NW, NW or SE routes from their breeding grounds. SV circled in black in the SE panel was also identified in our genome-wide association analyses (Fig. 4). The 5 % cutoff used for enrichment analyses is shown with a dotted line for each phenotype.

One GO term was enriched in the SE migrants: ‘MAPK cascade’ (mitogen-activated protein kinase cascade). This cascade is highly conserved across vertebrates and important for translating extracellular signals to intracellular responses. Of particular relevance to seasonal migration, MAPK cascades facilitate learning and memory, consolidating learning following specific behaviors and eliciting memory formation (51–53). MAPK cascades are also important for mounting immune responses in many vertebrates. Considerable research has focused on the relationship between immunity and migration, with several studies suggesting that migrants suppress their immune system on migration, allowing them to allocate more resources to this costly life history event. It has also been noted that migrants with different routes and wintering sites likely encounter different parasites throughout their annual cycle; local adaptation associated with this variation may also drive changes in the immune system (54–56).

### Genome-wide association analyses and individual migratory traits

The former analyses used local estimates of differentiation to identify genomic regions that distinguish the main migratory phenotypes present in blackcaps. In this last set of analyses, we broke migration behavior down into distinct traits and examined the genetic basis of each one with GWAS. We used all birds in these analyses (i.e., added birds exhibiting intermediate [S] orientations back in to the analysis along with those breeding in the Netherlands) and focused on seven traits: direction (the vector orientation between breeding and wintering sites), distance (direct connection between breeding and wintering sites, in km), wintering location (longitude), and both the duration (days) and timing (date when birds reached the halfway point between breeding and wintering sites) of fall and spring migration.

We started our analyses by estimating PVE (the proportion of phenotype variation explained by genetic variation in our SV set) for each trait. We used Bayesian sparse linear mixed models (BSLMMs) for these analyses (57). BSLMMs use an MCMC algorithm to fit all variants to the phenotype simultaneously and control for population structure with a kinship matrix. We ran 20 million MCMC steps, extracting parameter values every 10,000 steps. Mean values of PVE across these steps ranged from 0.45 to 0.82 (direction = 0.73 ± 0.27 [SD], distance = 0.45 ± 0.29, winter longitude = 0.82 ± 0.28, fall timing = 0.47 ± 0.28, fall duration = 0.50 ± 0.30, spring timing = 0.77 ± 0.29, spring duration 0.46 ± 0.29). These values are relatively high, suggesting our SVs capture a good amount of the variation present in the migratory traits we measured. Standard deviations around these means, however, are quite wide and indicate we have limited precision in these estimates. Accordingly, we ran a complementary set of analyses obtaining polygenic scores (PGS) for each individual and trait. Specifically, we randomly masked phenotypic values for a subset of individuals and tested if we could predict these missing values with the remaining dataset. Migratory orientation was the only trait where predicted and actual phenotypic values were correlated (Fig. 4a; F_1,77_ = 3.92, p = 0.05, all p-values for remaining traits > 0.11). Combined, results from PVE and PGS analyses suggest that even though PVE values were relatively high for all traits, we can only be confident that our SVs capture sufficient variation in migratory direction. This does not necessarily mean there is no genetic basis to the remaining traits; rather, future work using larger sample sizes and additional variants (e.g., SNPs) may be needed to explain variation in these traits. For now, we focus our remaining analyses on migratory direction.

**Figure 4.**
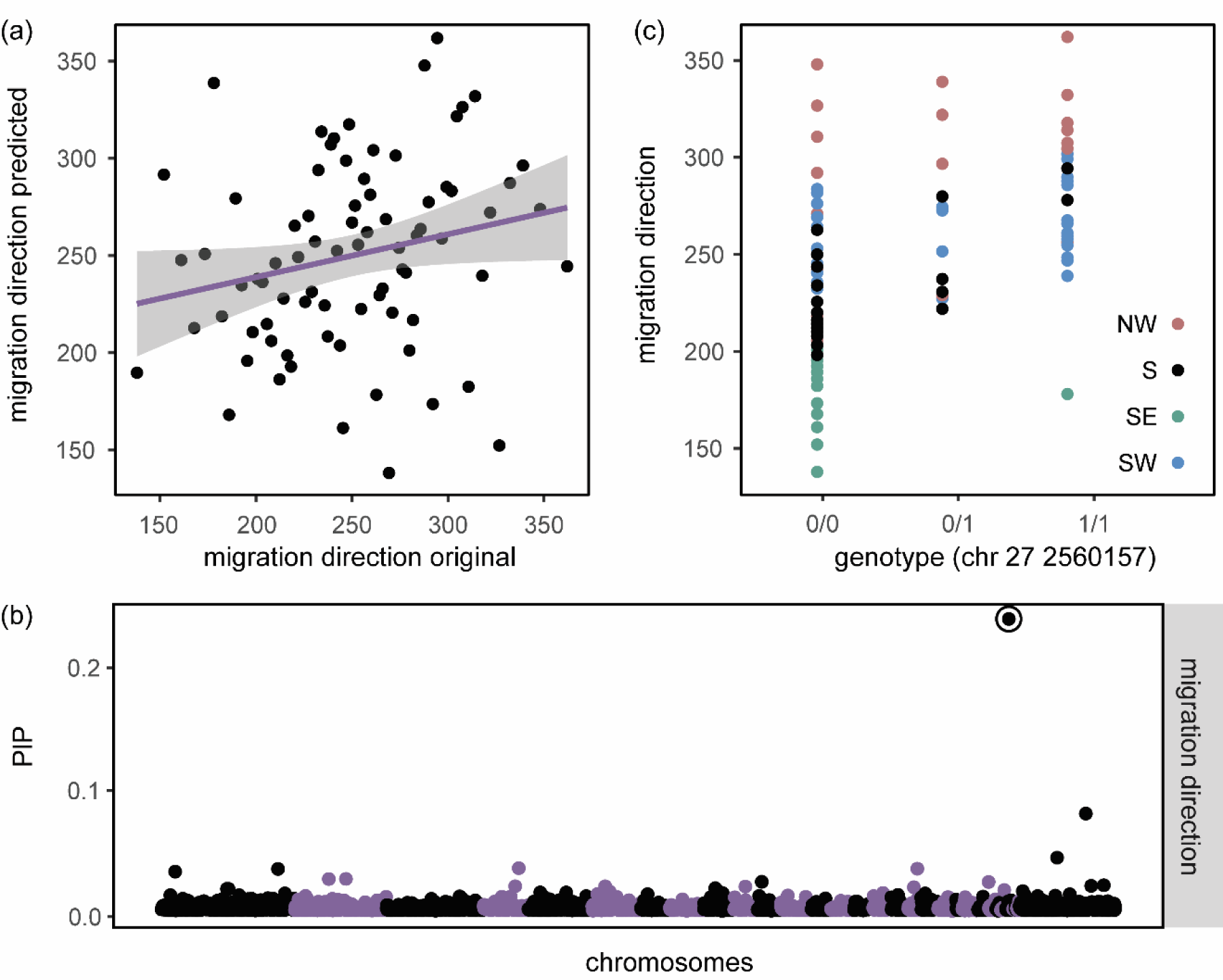
Results from genome-wide association analysis using migration direction during autumn. (a) Relationship between original values of migratory orientation and those predicted by cross-validation procedure (p=0.05). (b) Posterior inclusion probabilities (PIPs) for all variants, highlighting the deletion on chromosome 27 that exhibited elevated PBS in SE migrating birds (also highlighted in Fig. 3). (c) Relationship between genotypes at the deletion on chromosome 27 and migration direction (p<0.0001). Individuals are colored by their migratory phenotype.

Beyond PVEs, BSLMMs also estimate posterior inclusion probabilities (PIPs) for each variant. PIPs represent the proportion of MCMC iterations where a variant has a non-zero effect size. One variant stood out against the rest in our BSLMM for migration direction – a 710 bp deletion on chromosome 27 that had a PIP of 0.28 (Fig. 4b). Birds homozygous for the reference allele oriented in directions that were further east (Fig 4c). This variant also stood out in our analysis of genomic differentiation, exhibiting an elevated estimate of PBS in SE migrants (Fig 3). SE migrants were nearly fixed for the reference allele at this locus. The relationship between migratory orientation and genotypes at this locus remained even when we limited the dataset to S migrants (the group with the most variation in migratory direction)(F_2,16_ = 8.84, p = 0.003) suggesting variation at this SV is related to orientation, not just fixation in SE migrants. Combined, these findings suggest that this variant is under selection in SE migrants and helps control migratory orientation across blackcaps. Concerning the actual identity/effect of this variant, it occurs in an intron of TRA (T cell receptor alpha chain). TRA helps T cells respond to specific antigens in their cellular environment (58) and this finding continues to support a connection between immune response and migration in blackcaps; recall, the functional category ‘MAPK cascade’ was enriched in SE migrating birds. This pathway plays a role in memory and learning and is also important for mounting immune responses in many vertebrates. Together, these findings add support for an important role of immunity in the context of migration behavior, e.g. related to the fact that birds using different migratory routes are challenged with different environments throughout their annual cycle, and local adaptation associated with this variation may facilitate changes in the immune system.

## Conclusion

We conducted one of the most extensive studies of SVs in natural populations to date, using *de novo* assembled genomes to genotype 79 individually-phenotyped birds at thousands of SVs. We found evidence for purifying selection on SVs, suggesting they have an overall deleterious effect on fitness also supporting previous work on SVs. We also documented considerable overlap between SVs and transposable elements, suggesting transposable elements comprise a large proportion of SVs and genetic variation in the genome. We did not find evidence for genome-wide population differentiation between blackcaps with different migratory strategies, but our individual-based phenotypic characterization indicates local genetic variation at SVs does account for a large proportion of the phenotypic variation observed in specific migratory traits.

Seasonal migration is a complex behavior that comprises many traits. SVs like inversions are strong candidates for capturing loci that underlie complex behaviors and evidence from other systems has connected inversions with migration (e.g., an inversion on chromosome 1 underlies migratory direction in willow warblers)(27,32). We did not make such a connection here; we only identified a small number of small inversions. LD was reduced in these inversions suggesting they are suppressing recombination, but they did not exhibit signatures of selection and were not linked to any of our focal migratory traits. Blackcaps only began to diverge recently (30,000 years ago (25)) and seasonal migration is highly dynamic in this species (e.g., the NW population only recently [in the last 70 years] started to growing in size)(35). Accordingly, it is possible that inversions will capture genetic variation underlying migratory behavior in the future, but that is not currently the case. Inversions are only one type of SV; deletions, insertions, as well as translocations or duplications can also drastically alter phenotypes with important evolutionary consequences, such as the text book example of industrial melanism i.e. the darker morph of the peppered moth (59), or as an example within the songbirds, plumage color divergence between hooded and carrion crows (*Corvus corone cornix, C.c. corone* respectively) has been linked to a LTR retrotransposon insertions in crows, where hooded crows are homozygous for the insertion (13).

Future work using additional long read technologies (e.g., PacBio HiFi and HiC) may uncover additional variants that could be connected to migration in the system but for now, we conclude that seasonal migration has a highly polygenic basis in blackcaps. Beyond the deletion on chromosome 27, none of the SVs identified here stood out in our analyses of selection or GWAS. We reported similar findings in a previous study using genome-wide SNP data. Interestingly, the Ebox regulatory motif identified in the NW migrants here was also identified in enrichment analyses with population averaged phenotypes based on SNP data and could reveal a mechanism through which multiple migratory traits could be controlled by a similar mechanism. The Ebox motif is bound by transcription factors that regulate circadian rhythms in birds. Circadian rhythms are important for migration (e.g., songbirds like blackcaps switch from diurnal to nocturnal behavior on migration and circadian rhythms likely entrain circannual rhythms which are important for migratory timing). Perhaps seasonal migration in blackcaps is regulated by a small number of transcription factors that affect expression at multiple genes. In a general context we see recurrent pathways and functional categories connected to migration in many systems, including immunity, circadian rhythm regulation, learning and memory. Although the actual genes under selection do not seem to match in an across species comparison, we would assume that the adaptation of the central pathways needs to be optimized to and constrained by the migratory niche of each species or population, i.e. each species adapt their phenotype in a specific way, fitting to its ecological demands, which could explain consistency in general regulatory pathways despite the apparent lack of commonly identified genes.

## Materials and Methods

### Sampling and phenotypic analysis

We included data from 79 blackcaps in the present study. A subset of these birds were captured using mist nets on the breeding grounds in Austria (n = 45) and the Netherlands (n = 16); the remaining birds were captured on the wintering grounds in the UK (n = 18; Table S1). We obtained blood samples from each bird and fitted them with light-level geolocators using leg-loop backpack harnesses. Light-level geolocators record light intensity data at specific time intervals. These data are stored until the devices are retrieved at which point light intensity data are converted to day length and time of local midday and used to estimate daily longitude and latitude (60).

We describe methods used to analyze light-level geolocator data in full in Delmore et al. (2020b). Of relevance for the present study, we categorized birds into four broad phenotypic classes (migrating NW, SW, SE and S in fall) using their wintering locations. For birds wintering north of 37.5° N, we considered those west of 5° E to be southwest (SW) migrants, those east of 20° E to be SE migrants and those between 5 and 20° E to have intermediate southerly (S) routes. For birds wintering south of 37.5° N, we used a cut-off of 0° instead of 5° E to distinguish SW from S because these longer routes require less of a westerly component to reach the same longitude.

We estimated migration direction and distance by fitting a rhumb line between their breeding and wintering sites. We estimated timing by identifying the shortest distance route (i.e. a great circle routes) between their breeding and wintering sites and determining the date when birds reached 50% of the way between these sites. Duration was estimated as the number of days it took each bird to travel from early (30 %) to late (70 %) migration stages and speed as migration distance divided by duration.

### Assemblies and variant calling

High molecular weight DNA was extracted from blood samples, 10X Chromium libraries constructed and sequenced using Illumina technology (150 bp, paired end) by Novogene (HK). The mean molecule length of resulting libraries was 32,657 bp (range 10,062 – 54,206 bp) and sequencing reached a mean coverage of 55X (range 7 – 74X).

We called SVs using three pipelines. The first two pipelines relied on *de novo* assemblies of reads. Specifically, we assembled reads into two parallel pseudohaplotypes (phased contigs and scaffolds) with Supernova (61) and used two different approaches to align these pseudohaplotypes to the blackcap reference genome (62) and call genotypes in relation to the reference genome: (1) MUMmer4 (63) for alignment and MUM&Co (-g 1080000000 -b)(6) to call genotypes and (2) Minimap2 (64) for alignment and SVIM-Asm (diploid, tandem duplications as insertions and interspersed duplications as insertions) (65) for genotype calling. We aligned 10X reads directly to the blackcap reference for the third pipeline. We used LongRanger wgs for alignment (average mapping rate of 87%) and the same program to genotype SVs (--vcfmode gatk)(66).

Once SVs were genotyped, we used a series of filters to identify high quality variants. Starting at the level of individuals, we limited the dataset to variants that had been identified by at least two callers and had matching genotypes. We used SURVIVOR (1000 2 0 0 0 50) (67) and a custom R script to conduct this filtering. Variants with strings of >10Ns were also removed to reduce potential errors caused by contig scaffolding. We generated a multi-individual vcf (i.e., merged variants across individuals) using SURVIVOR (1000 4 0 0 0 50) and limited our analyses to SVs with maf > 0.05 using vcftools (68). We focused on five types of SVs: insertions (sequence inserted into query), deletions (sequence deleted from the query), tandem duplications (sequence duplicated in the query), inversions (sequence with reversed orientation) and translocations (sequence moved between chromosomes).

Following Hamala et al. (6), we chose a random set of 50 SVs for visual validation in IGV. We used alignments from LongRanger and confirmed the presence of all but one of these SVs (see examples in Fig. S2, including one of the main variants identified in our subsequent analyses), suggesting false positives are rare in our dataset and likely related to the stringent filtering we applied.

### Population genetics and GWAS

We used AFS to examine the overall fitness effects of SVs. We constructed these AFS using vcf2sfs in R (69). We used scripts from Hämälä et al. (2021) to estimate linkage disequilibrium (squared Pearson’s correlation coefficients) between arrangements at inversions and a random set of colinear regions with the same distribution of sizes as inversions.

We used a PCA to examine genome-wide patterns of genomic differentiation. This analysis was conducted using smartpca (EIGENSOFT version 5.0).

We used estimates of PBS to identify SVs exhibiting signatures of selection. PBS is similar to F_ST_ but can be used with more than two populations and identifies selection specific to one population. We estimated this parameter in two steps, calculating F_ST_ between NW, SW and SE migrants using the estimator derived by Hudson (70) and scripts from (6). These estimates of F_ST_ were then converted to PBS following (71)(T = log transformed estimates of F_ST_, example is for SE population): (T^SE-NW^ + T^SE-SW^ – T^NWSW^)/2.

We used two different programs to look for functional enrichment at genes overlapping SVs showing evidence for selection (i.e., with PBS values in the top 5% of the distribution): (1) BINGO (72) to look for enrichment of specific GO categories and go:Profiler (73) to look for enrichment in additional functional databases, including biological pathways, regulatory motifs of transcription factors and microRNAs and protein-protein interactions. We used a custom annotation for the blackcap in BINGO and an annotation for the chicken in go:Profiler.

GWAS were run as BSLMMs in GEMMA (57). BSLMMs are adaptive models that include linear mixed models (LMM) and Bayesian variable selection regression (BVSR) as special cases and that learn the genetic architecture from the data. These models are run separately for each phenotype but allele frequencies at all variants are considered together and included as the predictor variable. A kinship matrix is also included to control for factors that influence the phenotype and are correlated with genotypes (e.g., population structure). We ran four independent chains for each BSLMM, with a burn-in of 5 million steps and a subsequent 20 million MCMC steps (sampling every 1000 steps). We report one hyperparameter from this model (PVE: the proportion of variance in phenotypes explained by all SVs, also called chip heritability) and focus on two variant specific parameters: PIP (posterior inclusion probabilities) and (β, variant effects). We calculated genetic correlations between traits by identifying SNPs with PIP > 0.01 and correlated model averaged estimates of β (β weighted by their PIPs) (74,75). In order to facilitate comparisons across traits and limit the effects of outliers, we normal quantile-transformed all of our phenotypic traits before running these analyses. We also regressed each trait against sex to remove the effects of this variable on migratory traits.

We used a used a cross-validation procedure to obtain polygenic scores for each individual (and trait)(75,76). Specifically, we masked the phenotype of 25 % of the sampled individuals and reran BSLMMs using the remaining individuals and same parameters as the original BSLMM (with only one MCMC chain and the ‘predict −1’ plugin in GEMMA). We repeated this procedure four times for each trait, obtaining predicted values (polygenic scores) for each individual and used linear model to estimate correlation between predicted values and the original phenotype of each bird.

## Acknowledgments

We thank Matthias Weissensteiner and Claire Mérot for advice early in our analysis; Tuomas Hämäla and Hannah Justen for help implementing scripts for a subset of our population genetic and GWAS analyses, respectively; and Corinna Langebrake, Georg Manthey and Joe Wynn for general discussion. This work would not be possible without the enthusiasm and support received in the field to collect both accurate phenotype data as well as genetic material. We are particularly grateful to Ben Sheldon, Robbie Phillips, Greg Conway, Graham Roberts, Tania Garrido-Garduño, Britta Meyer, Timo Hasselmann, Hannah Justen, Juan Sebastian Lugo Ramos, Ivan Maggini, Wolfgang Vogl, Leonida Fusani next to many more keen fieldworkers and houseowners for assistance generating the phenotypic dataset. All work was carried out under approval of the respective institutional ethics and animal welfare committee and national authorities. Specifically, permit numbers for work in Austria: GZ BMWFW-68.205/0048-WF/V/3b/2016 and BMWFW-68.205/0139-WF/V/3b/2016 according to §§ 26ff. of Animal Experiments Act, TVG 2012.In the UK, geolocator deployment was approved by the University of Oxford Animal Welfare Ethical Review Body, and fieldwork was conducted under licenses from the British Trust for Ornithology, approved by the Special Methods Technical Panel. Permit number for work in the Netherlands: AVD801002016519 valid 27-6-2016 through 31-5-2021, issued by the Centrale Commissie Dierproeven. This project was supported by funding from the Max Planck Society (MFFALIMN0001 grant to ML), the German Science Foundation (project Nav05 within SFB 1372 – Magnetoreception and Navigation in Vertebrates to ML) and the National Science Foundation (National Science Foundation Grant IOS-2143004 to KED).

## Author contributions

Conceptualization: K.E.D., M.L.; sampling: K.E.D., B.M.V.D., T.C., H.J., M.L.; formal analysis: K.E.D., K.U.; writing: K.E.D. with input from B.M.V.D., K.U., H.J., M.L.

